# Premovement inhibition protects motor actions from interference

**DOI:** 10.1101/2021.04.26.441384

**Authors:** Aaron N. McInnes, Ottmar V. Lipp, James R. Tresilian, Ann-Maree Vallence, Welber Marinovic

## Abstract

Shortly before movement initiation, the corticospinal system undergoes a transient suppression. This phenomenon has been observed across a range of motor tasks, suggesting that it may be a obligatory component of movement preparation. We probed whether this was also the case when the urgency to perform a motor action was high, in a situation where little time was available to engage in preparatory processes. We controlled the urgency of an impending motor action by increasing or decreasing the foreperiod duration in an anticipatory timing task. Transcranial magnetic stimulation (TMS; experiment one) or a loud acoustic stimulus (LAS; experiment two) were used to examine how corticospinal and subcortical excitability were modulated during motor preparation. Preparatory inhibition of the corticospinal tract was absent when movement urgency was high, though motor actions were initiated on time. In contrast, subcortical circuits were progressively inhibited as the time to prepare increased. Interestingly, movement force and vigour were reduced by both TMS and the LAS when movement urgency was high, and enhanced when movement urgency was low. Our findings indicate that preparatory inhibition may not be a obligatory component of motor preparation. The behavioural effects we observed in the absence of preparatory inhibition were induced by both TMS and the LAS, suggesting that accessory sensory stimulation may disrupt motor output when such stimulation is presented in the absence of preparatory inhibition. We conclude that preparatory inhibition may be an adaptive strategy which can serve to protect the prepared motor action from external interference.

---

During preparation for a motor action, the corticospinal (CS) tract goes through systematic changes of excitability. Measurement of CS excitability via transcranial magnetic stimulation (TMS) of the primary motor cortex (M1) approximately 100 ms before electromyogram (EMG) onset in the effector muscle indicates an increase in excitation of CS neurons (Chen et al., 1998; Leocani et al., 2000; Starr et al., 1988). However, prior to this ramp-up of excitability, there is a period of CS inhibition, demonstrated by a gradual suppression of motor evoked potentials (MEPs) induced by TMS, up until the point excitation occurs (Hasbroucq et al., 1997a; Hasbroucq et al., 1997b; Hasbroucq et al., 1999; Ibáñez et al., 2020; Marinovic et al., 2011). There are three main explanations which have been put foward regarding the role of this suppression of motor circuits before movement onset. First, the competition resolution account suggests that CS inhibition may be necessary to suppress the initiation of competing response selections (Burle et al., 2004). Second, the impulse control hypothesis proposes that inhibition may be necessary to prevent a prepared movement from being triggered prematurely (Duque et al., 2010, 2017; Duque & Ivry, 2009). Finally, the spotlight hypothesis suggests premovement inhibition allows the speeded initiation of movement by increasing the signal to noise ratio in motor circuits (Greenhouse et al., 2015).

The available data, however, are not completely consistent with these three explanations. In choice RT time tasks, CS inhibition has been found to increase in the selected effector, rather than the non-selected effector after an action is specified (Duque & Ivry, 2009). This finding is not consistent with the competition-resolution hypothesis. Furthermore, the impulse control explanation implies that any facilitatory input to the CS tract should be suppressed. This is incompatible with findings that there is specificity in the suppression of volleys evoked by TMS (Hannah et al., 2018). In addition, shorter RTs are associated with greater levels of preparatory inhibition (Hannah et al., 2018), and CS suppression is observed in self-timed actions which do not rely on initiation taking place at a particular time (Ibáñez et al., 2020), findings of which are incompatible with preparatory inhibition acting to prevent premature initiation. Finally, the spotlight account is incompatible with the observation of CS suppression in non-selected muscles – it could be argued that this would be maladaptive in that it may lead to an increased risk of unintentionally triggering task-irrelevant actions. As such, it is unclear whether preparatory inhibition serves an entirely different role, or whether the phenomenon involves the combination of some or all of the above explanations (Duque et al., 2014). For example, a more global inhibition may initially take place at a short-time scale, acting on non-selected effectors to prevent their unintentional triggering. This may eventually unfold into a more specific inhibition acting in selected effectors consistent with the impulse control and/or spotlight hypotheses.

Preparatory inhibition has been observed using different neurophysiological techniques besides TMS. These include the Hoffman reflex (Derosiere, 2018; Hannah et al., 2018) and the eye-blink reflex, which, similarly to the CS tract, has been shown to undergo a facilitation close to movement onset (Lipp et al., 2001; Marinovic et al., 2013) and suppression earlier in preparation (Anthony & Putnam, 1985; Nguyen et al., 2020). While the Hoffman reflex can be exploited to derive information regarding the excitability of separate populations of corticospinal neurons, the startle-blink reflex can be used as a measure of global motor-related subcortical excitability (Kumru & Valls-Solé, 2006). One advantage of using startling stimuli to probe the excitability of subcortical circuits is that it also has a well-known effect on response initiation and execution: the StartReact effect (Anzak et al., 2011; Marinovic et al., 2016; McInnes et al., 2020; Valls-Solé et al., 1999). More specifically, by employing startling stimuli, one is able to test both the excitability of subcortical circuits as well as the effects of the startle eliciting sounds on motor output (Kumru & Valls-Solé, 2006; Marinovic et al., 2013; Nguyen et al., 2020).

We hypothesised that time constraints imposed on preparation would limit the ability of motor circuits to undergo suppression during preparation. As such, we modified the urgency of an anticipatory action by shortening or lengthening the duration of time between the point at which an action should be prepared and when it should be initiated. Changes in CS and startle circuit excitability that occur shortly prior to action initiation were probed using TMS and a startling stimulus in experiments one and two, respectively. We predicted reduced CS inhibition when urgency is high and, due to the lack of sufficient inhibition, movements that are triggered by an intense sensory stimulus would occur earlier and with greater vigour than movements for which ample time was allowed for preparation. Furthermore, if a generalised suppression of motor circuitry precedes a more specific one, then time-related constraints on preparation should act to a lesser extent on the suppression of subcortical circuits (potentially representing global inhibition) than on the more focal inhibition which would be observed in the CS tract. In contrast, when sufficient time for motor preparation is allowed, we expected greater CS and subcortical inhibition and smaller behavioural effects induced by accessory stimulation — smaller increase in vigour and force — than in the high urgency condition.

## Method – Experiment One

### Participants

Eighteen participants were recruited for experiment one (10 female; mean age = 25.33, SD = 7.71). All participants had normal or corrected vision and no apparent or known auditory impairments, neurological conditions, or injuries which may have impaired their ability to complete the task. Sixteen of the participants were self reportedly right handed and two participants reported being ambidextrous. Participants were screened for potential contraindications to TMS in accordance with the guidelines proposed by Rossi et al. (2009).

### Procedures

Participants were seated comfortably ∼70 cm in front of a 24.5” monitor (ASUS ROG PG258Q ; 240Hz refresh rate, 1920 x 1080 resolution). The experiment routines were run using custom scripts run in MATLAB 2015b. Timing and presentation of visual and electromagnetic stimuli were controlled using Psychtoolbox (v3.0.11) and MAGIC (v0.2; Habibollahi Saatlou et al., 2018) toolboxes. During the task, participants applied pressure to a force sensor (SingleTact 10 N calibrated sensor), which was held in a custom-made housing, with their right index finger in synchrony with the sweeping of a clock hand which was presented on the monitor. Participants were instructed to apply ballistic force with the index finger in abduction of the first dorsal interosseous (FDI) and to initiate the response in time with the clock hand reaching the 12 o’clock position. The first 10 trials in each experimental block were control trials in which no TMS was delivered. These initial 10 trials were excluded from analysis and were included in the experimental session only to ensure participants were familiar with the timing of that particular block before TMS was delivered. Participants completed two experimental blocks of 100 trials each (200 experimental trials total). In each trial, prior to the clock sweeping, the clock remained stationary on screen for a resting period of a duration randomised from a uniform distribution of 3 – 5 s, in order to prevent anticipatory motor preparation. The two speeds of the sweeping of the clock hand (high/low urgency) were varied between experimental blocks with block order counterbalanced across participants. The time from the start of the sweep to the intersection of the hand at the 12 o’clock position was either 350ms (high urgency), or 1400ms (low urgency). As the clock moved toward the 12 o’clock position, the centre remained white up until -25 ms prior to the intersection at 12 o’clock, and turned green for a duration of 50 ms so that the centre flashed green ± 25 ms around the expected time of movement onset. In 40% of trials, TMS was presented either 2 s after the beginning of the resting period during which the clock was stationary (baseline TMS), or 250 ms prior to the expected time of movement onset (probe TMS). Feedback regarding the temporal error of movement onset was provided at the end of each trial, with the exception of probe trials. Feedback was not provided in probe trials to avoid participants changing their responses due to the probe stimulus interfering with their ability to initiate their movements on time. For movements initiated within ± 25 ms of the clock hand intersection, a “Good timing!” message was presented in green text. For movements initiated < -25 ms or > 25 ms, a message of “Too early” or “Too late”, respectively, was presented in red text. Feedback regarding the temporal error of movement onset was also presented visually in a horizontal bar which depicted temporal error = 0, temporal error = 25, and temporal error = -25, along with temporal error of the current trial (see Figure 1). This feedback was presented for two seconds. Along with temporal error feedback, points were awarded to participants for each “good timing” response to encourage participants to initiate actions as close to the expected time of movement onset as possible. The ongoing score was presented along with temporal error feedback at the end of each trial. Prior to commencing experimental trials, participants completed 12 practice trials (six trials for each foreperiod length) and before beginning the task 20 TMS pulses were presented in order to measure MEPs at rest. Participants remained with their hands at rest in the same position they would hold the force sensor during the task and looked at a fixation cross presented on screen while the resting TMS pulses were presented.

**Figure 1.**
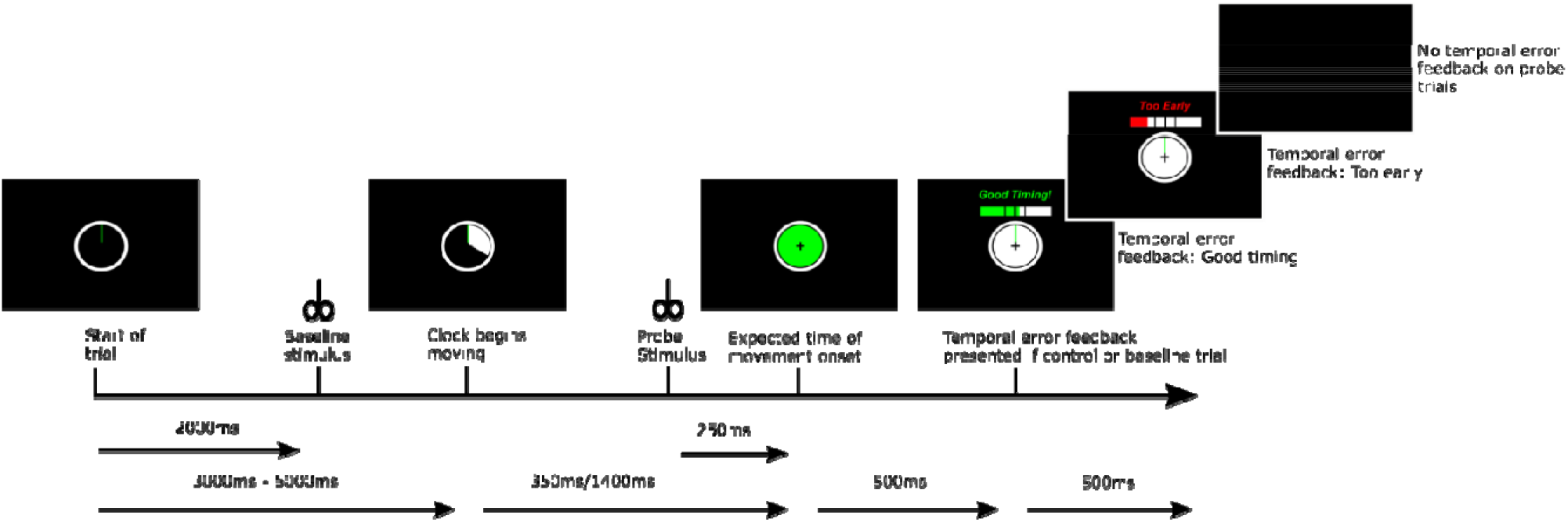
Sequence of events during experiment one. Note baseline and probe stimuli were presented on separate trials.

### Transcranial magnetic stimulation

TMS was delivered using a Magstim BiStim^2^ magnetic stimulator using a 70mm Magstim D70^2^ figure-of-eight coil. The coil was held tangentially to the participants’ scalp over the primary motor cortex (M1) and with the handle facing the rear of the head, placed at a 45° angle to the sagittal plane. Before commencing the experimental trials, the hotspot for the FDI (the primary agonist muscle) was located and the resting motor threshold was determined by using the Rossini-Rothwell procedure (Rossini et al., 1994) to find the lowest stimulus intensity to the nearest 1% of maximum stimulator output that could elicit a motor evoked potential with a peak to peak amplitude greater than 50 μV in five out of 10 test trials. Test pulses during the experiment were delivered at 120% of resting motor threshold. The mean resting motor threshold was 38.33% of maximum stimulator output (range = 32% – 49%). TMS was presented in 40% of trials (20% baseline timing, 20% probe timing) and trials were pseudorandomised so that no two consecutive trials could occur as a TMS trial.

### Data acquisition, reduction, and analysis

#### Acquisition of force and electromyogram data

Data were acquired using a National Instruments USB-6229 data acquisition device and sampled continuously at 2 kHz each trial. The data acquired from the force sensor were detrended and multiplied by a factor of 10 to convert the voltage output of the Singletact sensor to Newtons (N). These data were then used to determine temporal error of movement onset (difference in ms between the intersection of the clock hand at 12 o’clock and the time of movement onset, calculated using the algorithm suggested by Teasdale et al. (1993)), peak force (maximum force applied to the sensor over the course of a trial), and peak rate of force development (maximum derivative of the force signal over time occurring over the course of a trial; N/s). EMGs were recorded from the right FDI using bipolar 24 mm electrodes with a reference electrode placed over the styloid process of the right ulna. The EMG signal was amplified with a gain of x1000 using a pre-amplifier (Digitimer NeuroLog NL844) and amplifier (Digitimer NeuroLog NL820A) and the signal was band-pass filtered using a low-pass filter at 500 Hz (Digitimer Neurolog NL135) and high-pass filter at 20 Hz (Digitimer Neurolog NL144).

#### Processing of electromyogram data

All analyses were conducted using R software (v3.5.1). EMG data from FDI were downsampled to 1 kHz and EMG peak to peak amplitudes between 20 ms – 80 ms after TMS presentation were automatically calculated. All trials were visually inspected and peak to peak amplitudes were manually marked if they were incorrectly marked by the algorithm. Trials were excluded from analysis if visual inspection indicated significant noise, artifacts, or voluntary contraction, which obscured the detection of peak to peak amplitude, were present in the EMG record. Visual inspection was conducted blindly with respect to experimental condition. The manual rejection of trials resulted in the removal of 135 (participant median = 5, range = 0 – 21) trials. In addition, after manual trial rejection we calculated the root mean square of FDI EMG activity 200 ms prior to TMS presentation for all trials. Median root mean square of FDI EMG activity 200 ms prior to TMS presentation was calculated for each participant and if for a single participant any trial exceeded that median value by a factor of 1.4, the trial was excluded from the analysis of FDI EMGs. This resulted in the removal of a further 106 trials (participant median = 2, range = 0 – 25; n baseline timing = 58, n probe timing = 48). In total, 241 trials out of 1440 probe trials (16.74%) were excluded from the analysis of FDI EMGs. The findings observed from our analyses using these trial exclusion criteria were consistent with those obtained from analyses for which all trials were retained. In addition, we ran linear mixed-effects models to examine whether voluntary muscle contraction, as indicated by FDI EMG root mean square values 200 ms prior to TMS presentation, systematically differed as a function of foreperiod length, trial type, or as an interaction between the two.

#### Analysis of behavioural and electromyogram data

The behavioural data (temporal error of movement onset, peak force, peak rate of force development), and EMG data (MEP peak to peak amplitude) were subject to statistical analyses. In the analysis of behavioural data, control trials were excluded from analysis if their temporal error of movement onset was < -150 ms or > 150 ms. This resulted in the removal of 98 trials (participant median = 4, range = 0 – 29, 4.54% of all control trials) from the analysis of behavioural data. A series of linear mixed-effect models were conducted using the *lmer* function (lmerTest package, v3.0). Models were conducted with foreperiod length (350 ms, 1400 ms) and trial type (control, baseline, probe) as fixed factors and participant IDs as random factors. All valid trials were run in the models and all main effects and interactions were tested. The same models were conducted with MEP peak to peak amplitude as the dependent variable. In addition, EMG amplitudes of probe trials were calculated as a percentage of EMG amplitude in baseline trials to determine the magnitude of change in amplitude that occurs from baseline to probe responses. This involved calculating the median value of baseline trials for each foreperiod length and dividing each probe trial value by the median baseline value for its corresponding foreperiod length, and multiplying the result by a factor of 100 (i.e. 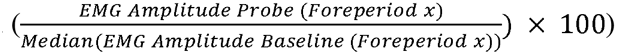. We also examined MEPs to the pre-experimental resting TMS, and compared these to MEPs to the experimental baseline TMS to examine whether CS excitability, as measured by MEPs in experimental baseline trials, was modulated by our manipulation of foreperiod length. Similarly to our calculation of amplitudes of EMG in probe trials as a percentage of amplitudes in baseline trials, we calculated MEP amplitude of experimental baseline trials as a percentage of median EMG amplitude of pre-experimental MEPs. The resulting *F* values, with Kenward-Rogers approximation for degrees of freedom, and *p* values from all linear mixed models are reported along with *R*^*2*^ values which were calculated using the *r2beta* function (r2glmm package, v0.1.2) in order to provide an estimate of effect size. Post-hoc tests were conducted using the *emmeans* function (emmeans package, v1.3.0) with the false discovery rate correction method for multiple comparisons (Benjamini & Hochberg, 1995). In addition to our linear mixed-effects analyses, we complemented our frequentist methods with Bayesian analyses using the *BayesFactor* package (v0.9.12) in order to provide the degree of support for the null hypothesis when such evidence would be relevant to the study aims. As such, BF_01_ values are reported to indicate the level of evidence to support the null hypothesis, and BF_10_ values are reported to indicate the level of evidence to support the alternative hypothesis. Furthermore, the *ttestBF* function was used to derive a Bayes factor (BF) indicating the degree of support the data provide for a facilitation of probe MEPs (> 100% of baseline), as well as a BF indicating the degree of support for an inhibition of probe MEPs (< 100% of baseline). The resulting BFs of a facilitatory effect were then divided by BFs indicating the probability of an inhibitory effect ([probability of data if excitatory effect / probability of data if null effect] / [probability of data if inhibitory effect / probability of data if null effect]). In this analysis, BF_10_ values are reported to indicate the level of support for a facilitatory effect and BF_01_ values are provided to indicate the level of support for an inhibitory effect.

## Results – Experiment One

### Urgency effects on temporal error

Average temporal error did not significantly differ between foreperiod lengths, as indicated by a non-significant main effect of foreperiod duration in the linear mixed-effects model of temporal error of movement onset, *F*_*(1*, 3446.3)_ = 1.16, *p* = .282, *R*^*2*^ = .000. The main effect of trial type was statistically significant, *F*_*(2*, 3446.3)_ = 11.76, *p* < .001, *R*^*2*^ = .007. Post-hoc tests indicated temporal error of movement onset was earliest for the probe TMS condition (*M* = - 14.71 ms, *SD* = 96.99), with later temporal error of movement onset occurring for the control condition (*M* = -7.83 ms, *SD* = 51.4; *p* = .003), and baseline TMS condition (*M* = -1.11 ms, *SD* = 56.8; *p* < .001). The interaction of trial type with foreperiod length was not statistically significant, *F*_*(2*, 3446.2)_ = 2.06, *p* = .128, *R*^*2*^ = .001. Figure 2A shows mean temporal error of movement onset for each trial type across both of the foreperiod durations.

**Figure 2.**
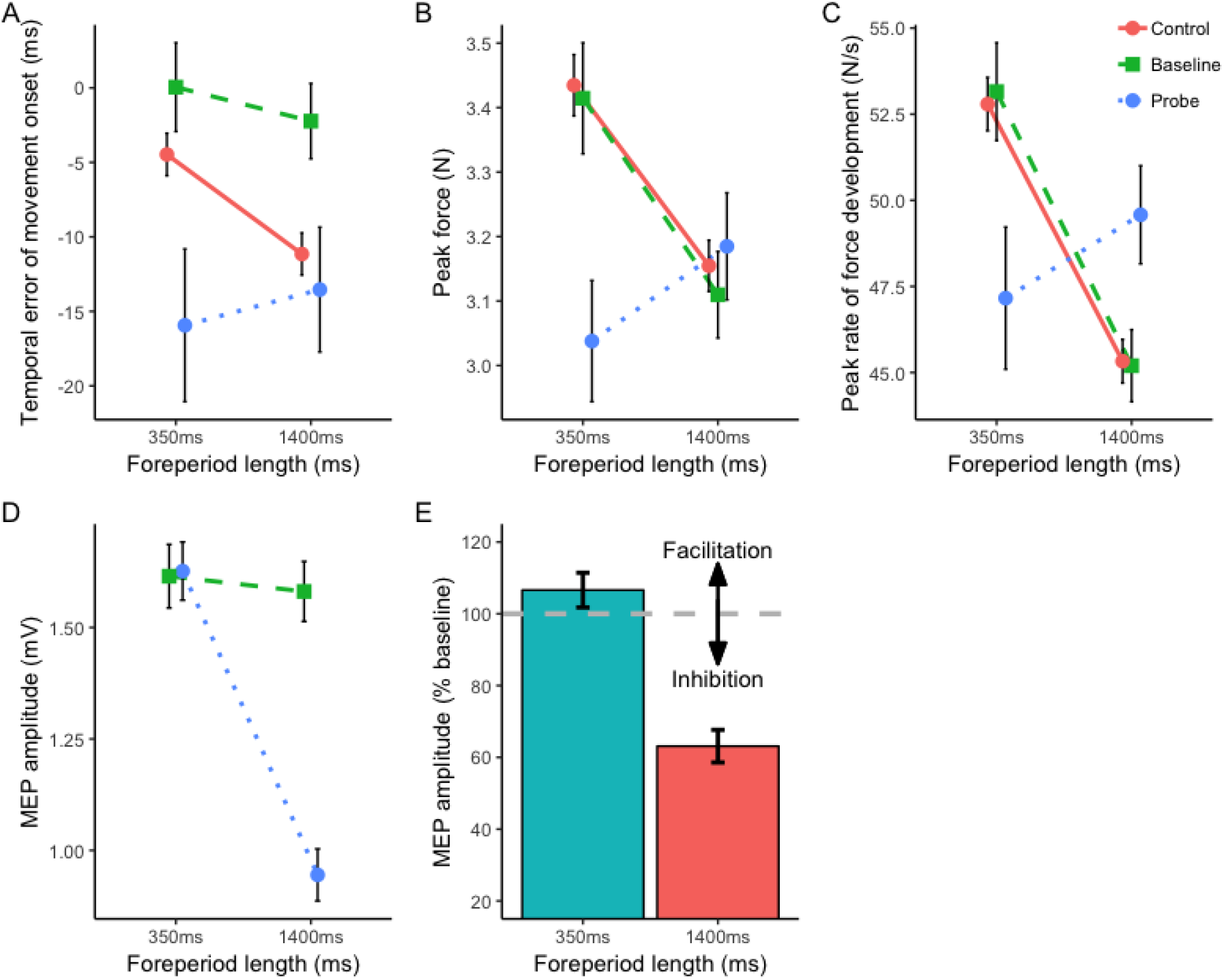
A). Temporal error of movement onset for control, baseline and probe trials over foreperiod lengths. B). Mean peak force of movements executed in control, baseline, and probe trials over foreperiod lengths. C). Mean peak rate of force development of movements executed in control, baseline, and probe trials over foreperiod lengths. D). Mean motor evoked potential (MEP) amplitudes for baseline and probe trials across each foreperiod length. E). MEP amplitudes in probe trials as a percentage of baseline amplitude over foreperiod lengths. Error bars represent standard error of the mean.

### Urgency effects on peak force and vigour

Analysis of peak force indicated that on average, responses in the 350 ms foreperiod duration (*M* = 3.35 N, *SD* = 2.07) were executed with greater force than those in the 1400 ms foreperiod duration (*M* = 3.15 N, *SD* = 1.73), as indicated by a significant main effect of foreperiod duration in the linear mixed model of peak force data, *F*_(1, 3479)_ = 7.94, *p* = .004, *R*^*2*^ = .002. The main effect of trial type was also statistically significant, *F*_(2, 3479)_ = 4.97, *p* = .006, *R*^*2*^ = .003. Average peak force was significantly reduced in probe TMS trials (*M* = 3.11 N, *SD* = 1.88), from control trials (*M* = 3.29 N, *SD* = 1.59; *p* = .005). Average peak force in baseline TMS trials (*M* = 3.26 N, *SD* = 1.61) was not significantly different from control trials (*p* = .583), nor probe TMS trials (*p* = .054). The linear mixed-effects model also showed a significant interaction of foreperiod length with trial type, *F*_(2, 3479)_ = 7.38, *p* < .001, *R*^*2*^ = .004. Post-hoc tests indicated a significant difference in peak force between control trials and probe TMS trials for the 350 ms foreperiod duration (*p* < .001), but not for the 1400 ms foreperiod duration (*p* = .459; see Figure 2B).

Analysis of peak rate of force development indicated that on average, responses in the 350 ms foreperiod length (*M* = 51.71 N/s, *SD* = 37.11) were executed with greater peak rate of force development than those in the 1400 ms foreperiod length (*M* = 46.17 N/s, *SD* = 28.24), as indicated by a significant main effect of foreperiod length in the linear mixed model, *F*_(1, 3479)_ = 23.51, *p* < .001, *R*^*2*^ = .007. The main effect of trial type was not statistically significant, *F*_(2, 3479)_ = 0.28, *p* = .754, *R*^*2*^ = .000. The linear mixed-effects model also showed a significant interaction of foreperiod length with trial type, *F*_(2, 3479)_ = 13.22, *p* < .001, *R*^*2*^ = .008. Post-hoc tests indicated a significant decrease in peak rate of force development from control trials to probe TMS trials for the 350 ms foreperiod length (*p* = .001), and in contrast, peak rate of force development showed a significant increase from control trials to probe trials during the 1400 ms foreperiod length (*p* = .012; see Figure 2C).

### Suppression of motor evoked potentials

Background EMG activity was assessed by analysing the root mean square of EMG 200 ms prior to the presentation of TMS. A linear mixed-effects model indicated a significant main effect of foreperiod length, *F*_(1, 1176.5)_ = 16.93, *p* < .001, *R*^*2*^ = .014 and indicated EMG background activity was greater in the 350 ms foreperiod length (*M* = 4.87 × 10^−3^ mV, *SD* = 1.63 × 10^−3^) than in the 1400 ms foreperiod length (*M* = 4.6 × 10^−3^ mV, *SD* = 1.61 × 10^−3^). However, this difference was small, with the difference between mean EMG root mean square in the 350 ms and 1400 ms conditions = 2.7 × 10^−4^ mV. Importantly, both the main effect of trial type, *F*_(1, 1176)_ = 0.79, *p* = .374, *R*^*2*^ = .001, and the interaction of foreperiod length with trial type, *F*_(1, 1176)_ = 0.33, *p* = .568, *R*^*2*^ = .000, were not statistically significant. A Bayesian linear model of the interaction of trial type with foreperiod length indicated BF_01_ = 346.12, providing decisive evidence for the null hypothesis (Jeffreys, 1961). Furthermore, we calculated the baseline MEP amplitude for each foreperiod duration condition as a percentage of median MEP amplitude when participants were at rest prior to the commencement of experimental trials. A linear model indicated a statistically significant main effect of the foreperiod duration condition of the block which the baseline probe was contained in, *F*_(1, 581.41)_ = 4.9, *p* = .027, *R*^*2*^ = .008. These percentages were larger for the 350 ms condition (*M* = 208.76%, *SD* = 233.51) than for the 1400 ms condition (*M* = 180.92%, *SD* = 196.32), suggesting there may have been some ongoing enhancement of corticospinal excitability prior to the start of the clock sweep for the 350 ms condition.

MEPs showed a modulation of amplitude with the manipulations of foreperiod length and TMS timing, with the linear mixed model of MEP amplitude showing statistically significant main effects of foreperiod length, *F*_(1, 1177.4)_ = 40.48, *p* < .001, *R*^*2*^ = .033, and trial type, *F*_(1, 1176.1)_ = 29.91, *p* < .001, *R*^*2*^ = .025. The interaction of foreperiod length and trial type was also statistically significant, *F*_(1, 1176.1)_ = 32.59, *p* < .001, *R*^*2*^ = .027. Post-hoc tests indicated that baseline MEP amplitudes did not differ between the foreperiod duration conditions (*p* = .717). Furthermore, for the 1400 ms foreperiod length, MEP amplitudes were significantly reduced relative to baseline TMS (*M* = 1.58 mV, *SD* = 1.16) in probe TMS trials (*M* = 0.94 mV, *SD* = 1.01; *p* < .001). This reduction in MEP amplitude from baseline (*M* = 1.61, *SD* = 1.23) in probe (*M* = 1.63, *SD* = 1.13) TMS trials was not present during the 350 ms foreperiod length (*p* = .864). A Bayesian paired-samples t-test provided no support (BF_01_ = 1.92 × 10^−3^) for the null hypothesis of differences in participant mean MEP amplitudes between baseline and probe trials for the 1400 ms condition, and provided decisive evidence (Jeffreys, 1961) for the alternative hypothesis, BF_10_ = 521.48. Conversely, for the 350 ms foreperiod length, a Bayesian paired-samples t-test provided substantial evidence for the null hypothesis of differences in MEP amplitudes between baseline and probe TMS trials, BF_01_ = 3.84 (Jeffreys, 1961). Figure 2D shows mean MEP amplitudes across probe and baseline TMS trials for each of the foreperiod lengths. MEP amplitudes in probe TMS trials were also calculated as a percentage of the median amplitude of MEPs in baseline TMS trials for each foreperiod length. A linear mixed model indicated a significant main effect of foreperiod length for these percentages, *F*_(1, 590.33)_ = 84.97, *p* < .001, *R*^*2*^ = .126. Probe MEP amplitudes as a percentage of baseline MEP amplitudes trended toward facilitation for the 350 ms foreperiod length, with mean percentage being > 100% (*M* = 106.6%, *SD* = 83.35). However, a one-sample t-test of the mean of these percentages against a mean of 100% was not statistically significant (*p* = .127). Probe MEP amplitudes showed a clear inhibition during the 1400 ms foreperiod length (*M* = 63.12%, *SD* = 78.7), and a one-sample t-test indicated these percentages were significantly reduced from 100% (*p* < .001). We calculated BFs which indicate the magnitude of evidence to suggest observation of a facilitatory effect in probe MEP amplitudes over baseline amplitudes, for each foreperiod duration. Trials in the 350 ms foreperiod duration indicated BF_10_ = 14.49, providing a strong evidence (Jeffreys, 1961) to suggest facilitation of probe MEPs. In contrast, analysis returned a BF_10_ = 2.61 x 10^−22^ for the 1400 ms condition, providing no support of facilitation in this condition, and rather providing decisive evidence for an inhibitory effect, BF_01_ = 3.83 x 10^21^. Figure 2E shows mean probe MEPs as a percentage of median baseline MEPs for each foreperiod duration.

## Method – Experiment Two

### Participants

Twenty-three healthy participants were recruited for experiment two (12 female; mean age = 23.61, *SD* = 5.84). Twenty-two participants were self-reportedly right-handed, and one participant reported being ambidextrous. All participants had normal or corrected vision and no apparent or known auditory impairments, neurological conditions, or injuries which may have impaired their ability to complete the task.

### Procedures

A similar procedure was employed in experiment two, with the following exceptions. Three foreperiod lengths were employed, high urgency (350 ms), medium urgency (700 ms), and low urgency (1400 ms), in order to reference a time point at which preparatory inhibition emerges. Each of the foreperiod durations were randomised to one of three blocks which consisted of 100 trials each (300 trials total). Furthermore, rather than single pulse TMS, a loud acoustic stimulus (LAS) was presented at the same timings as the TMS in experiment one. A LAS was used in this experiment as it has recently been shown that preparatory inhibition can be observed in the modulation of the startle eyeblink response (Nguyen et al., 2020), and it would allow us to discern whether the behavioural effects we observed in experiment one were due to activation of the corticospinal tract or whether to the accessory auditory effects of the TMS coil’s discharge. Prior to commencing experimental trials, participants completed 12 practice trials (4 trials for each foreperiod length). The LAS was also presented four times to participants before beginning the task in order to measure their orbicularis oculi (OOc) EMG response at rest.

### Data acquisition, reduction and analysis

EMG from the right OOc was recorded with surface bipolar 8 mm Ag/AgCl sintered electrodes with a 24 mm reference electrode placed over the right mastoid process. Onset latency and amplitude of OOc EMG in experiment two were measured using several steps. EMG data from OOc were downsampled to 1 kHz, rectified using the *rectification* function (biosignalEMG package, v2.1.0), and the *rollapply* function (zoo package v1.8) was used to smooth data using a five-point moving average. The latency of OOc EMG onset was detected using the Bonato (1998) method with the *onoff_Bonato* function (biosignalEMG package, v2.1.0; sigma n = two times the standard deviation of activity within 0 – 200 ms prior to the LAS). Multiple passes of the Bonato method were run until onset of EMG could be detected. If no onset of OOc EMG could be detected, the threshold was increased by an increment of 0.2x(Baseline variability) for a maximum of 10 passes, after which the threshold was decreased from 1 by increments of 0.2x(baseline variability) for a maximum of two passes, until an onset of EMG could be detected between 20 – 80 ms with respect to LAS onset.

Amplitude of the EMG was automatically calculated as the difference between the maximum of the rectified EMG signal after blink onset and baseline amplitude of the rectified signal. All trials were visually inspected, and adjustments were made to the latency of EMG onset or amplitude if necessary. Trials were excluded from analysis of OOc EMG if their onset was < 20 ms or > 80 ms (Blumenthal et al., 2005), if no EMG response occurred, or if excessive noise, artifacts, or voluntary activation within 20 ms of LAS onset were present in the EMG record. Seven participants were excluded from analysis of EMG data in experiment two due to excessive noise or artifacts in EMG recordings, or insufficient OOc EMG response to the LAS, and as such, EMG data are reported for the remaining 16 participants. After removal of these participants, our trial exclusion criteria resulted in the exclusion of 128 trials (9.27% of all LAS trials). Finally, T-scores were calculated for OOc EMG amplitude using the *rescale* function and setting *M* = 50 and *SD* = 10. Exclusion of control trials of behavioural data for which temporal error of movement onset was < -150 ms or > 150 ms resulted in the exclusion of 117 trials in experiment two (2.83% of all control trials).

### Loud acoustic stimulus

A LAS was presented in 40% of trials; 20% at baseline timing and 20% at probe timing. Trials were pseudorandomised to either control, baseline LAS, or probe LAS trials so that no two consecutive trials could occur as a LAS trial. The onboard audio of the computer used to run the experiment generated the LAS as brief bursts of white noise (50 ms burst duration with a rise and fall time < 1.5 ms). The LAS was presented binaurally through stereophonic active noise cancelling headphones (Bose QC25). At a distance of 2 cm from the speaker cone, the peak amplitude of the LAS was measured at 105 dBA.

## Results - Experiment two

### Urgency effects on temporal error of movement onset

On average, movement onset was shortest for the 700 ms foreperiod length (*M* = -15.64 ms, *SD* = 62.91), and the 1400 ms condition (*M* = -12.88 ms, *SD* = 58.59), with the 350 ms foreperiod length resulting in the longest movement onsets (*M* = 3.53, *SD* = 60.24). The linear mixed-effects model of temporal error of movement onset data indicated a statistically significant main effect of foreperiod duration, *F*_*(2*, 6750)_ = 50.7, *p* < .001, *R*^*2*^ = .015. Post-hoc tests indicated in comparison to the 350 ms condition, both the 700 ms (*p* < .001) and 1400 ms (*p* < .001) foreperiod durations resulted in significantly earlier movement onsets. The main effect of trial type was also statistically significant, *F*_*(2*, 6750.1)_ = 42.02, *p* < .001, *R*^*2*^ = .012, with movements on probe trials (*M* = -24.49 ms, *SD* = 88.05) being initiated earlier on average than on baseline (*M* = -4.5 ms, *SD* = 67.32, *p* < .001) and on control trials (*M* = -7.08 ms, *SD* = 57.18, *p* < .001), but with no significant difference between control and baseline trials (*p* = .267). The interaction of foreperiod length with trial type was also statistically significant, *F*_(4, 6750)_ = 2.76, *p* = .026, *R*^*2*^ = .002. However, analysis of the difference in temporal error between probe trials and control trials indicated the main effect of foreperiod duration was not statistically significant, *F*_(2, 665)_ = 0.18, *p* = .832, *R*^*2*^ = .001. Figure 3A shows mean temporal error of movement onset for control, baseline, and probe trials for each foreperiod duration.

**Figure 3.**
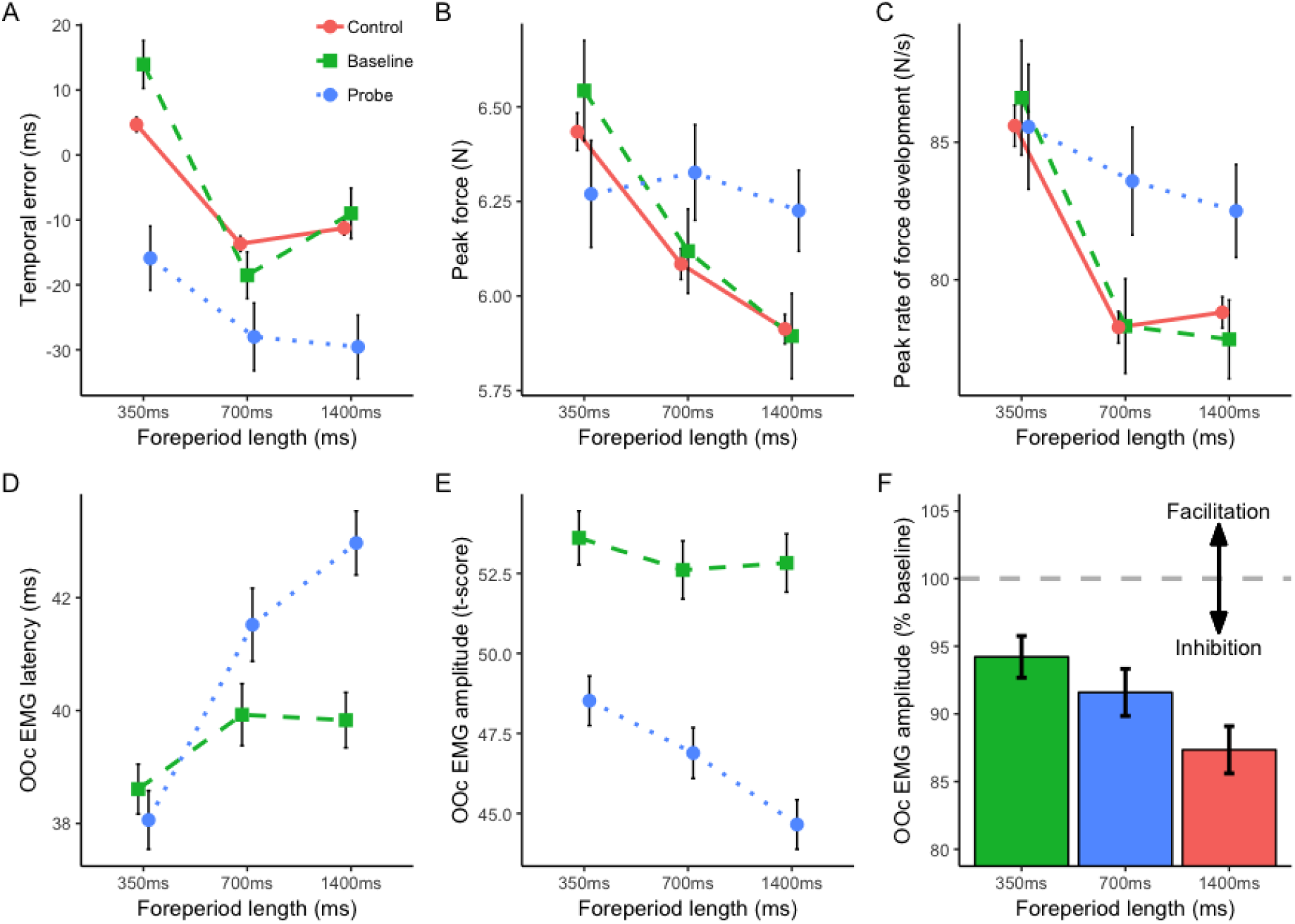
A). Mean temporal error of movement onset across control, baseline, and probe trials for each foreperiod length. B). Mean peak force of movements across control, baseline, and probe trials for each foreperiod length. C). Mean peak rate of force development of movements across control, baseline, and probe trials for each foreperiod length. D). Mean latency of orbicularis occuli electromyogram onset for baseline and probe trials across each foreperiod length. E). Mean t-scores of Orbicularis oculi electromyogram amplitude in baseline and probe trials over foreperiod lengths. F). Mean orbicularis oculi electromyogram amplitude after the probe stimulus as a percentage of median electromyogram amplitude after the baseline stimulus. Error bars represent standard error of the mean.

### Urgency effects on peak force and rate of force development

Mean peak force was greatest for the 350 ms foreperiod duration (*M* = 6.43 N, *SD* = 2.43), with lower forces executed in the 700 ms (*M* = 6.11 N, *SD* = 2.01) and 1400 ms (*M* = 5.94 N, *SD* = 1.92) conditions. The main effect of foreperiod duration was statistically significant for peak force, *F*_*(2*, 6752)_ = 13.27, *p* < .001, *R*^*2*^ = .008. Post-hoc tests indicated in comparison to the 350 ms condition, average peak force was significantly lower in both the 700 ms (*p* = .003) and 1400 ms (*p* < .001) foreperiod durations. The main effect of trial type was not statistically significant, *F*_*(2*, 6752)_ = 1.77, *p* = .17, *R*^*2*^ = .001, however, the interaction of foreperiod length with trial type on peak force was significant, *F*_*(4*, 6752)_ = 2.52, *p* = .039, *R*^*2*^ = .004. Post-hoc tests indicated a significant difference in peak force between control and probe trials for the 1400 ms foreperiod duration (*p* = .031), but not for the 700 ms (*p* = .071), or 350 ms foreperiods (*p* = .177). Figure 3B shows mean peak force executed during control, baseline, and probe trials for each foreperiod duration.

Mean peak rate of force development was also greatest in the 350 ms foreperiod length, (*M* = 85.7 N/s, *SD* = 37.05), with movements showing decreased peak rate of force development in the 700 ms (*M* = 78.82 N/s, *SD* = 29.13) and 1400 ms (*M* = 79.09 N/s, *SD* = 27.84) foreperiods. This was indicated by a significant main effect of foreperiod duration, *F*_*(2*, 6752)_ = 17.75, *p* < .001, *R*^*2*^ = .005. The main effect of trial type was also statistically significant for peak rate of force development, *F*_*(2*, 6752)_ = 4.13, *p* = .016, *R*^*2*^ = .001, with mean peak rate of force development being greatest in probe LAS trials (*M* = 83.88 N/s, *SD* = 34.77), with lower peak rate of force development executed in control trials (*M* = 80.9 N/s, *SD* = 31.5) and baseline LAS trials (*M* = 80.92 N/s, *SD* = 31.28). The interaction of trial type with foreperiod length was not statistically significant, *F*_*(4*, 6752)_ = 1.38, *p* = .237, *R*^*2*^ = .001. Figure 3C shows mean peak rate of force development executed during control, baseline, and probe trials for each foreperiod duration.

### Orbicularis oculi electromyogram onset latency and amplitude

The amplitudes of OOc EMG responses to the LAS at baseline timing for each foreperiod duration were calculated as a percentage of EMG amplitudes to the LAS presented at rest prior to experimental trials. These percentages (M_350_ = 107.86%, SD = 23.86; M_700_ = 106.48%, SD = 24.75; M_1400_ = 106.85%, SD = 23.45) were not significantly different between different foreperiod durations, *F*_*(2*, 393.5)_ = .152, *p* = .859, *R*^*2*^ = .001. This suggests motor preparation at the time the baseline LAS was presented was not significantly different between foreperiod durations. A Bayesian linear model was run to examine the degree of support for the null hypothesis and returned BF_01_ = 34.19, indicating very strong evidence (Jeffreys, 1961) for the null hypothesis of differences between foreperiod durations. We also examined OOc EMG amplitude in baseline trials and a Bayesian linear model provided very strong evidence to suggest EMG amplitude did not differ as a function of foreperiod duration, BF_01_ = 33.92. Similarly, a Bayesian linear model of OOc onset latency after the baseline LAS indicated BF_01_ = 14.50, suggesting strong evidence to support the null hypothesis of OOc latency being modulated by the foreperiod duration of experimental trials (Jeffreys, 1961). Statistically significant main effects of foreperiod duration, *F*_*(2*, 870.07)_ = 21.92, *p* < .001, *R*^*2*^ = .048, and trial type, *F*_*(1*, 870.25)_ = 12.01, *p* < .001, *R*^*2*^ = .014 were observed for the linear mixed-effects models of OOc EMG onset latency. Furthermore, the interaction of foreperiod length with trial type was statistically significant, *F*_*(2*, 870.1)_ = 7.27, *p* < .001, *R*^*2*^ = .016. Post-hoc tests indicated onset of EMG in probe trials (*M* = 38.06 ms, *SD* = 6.35) were not significantly different from baseline trials (*M* = 38.61 ms, *SD* = 5.55) for the 350 ms foreperiod length (*p* = .419). However, EMG onset latencies for probe trials were significantly delayed in the 700 ms (*M* = 41.52 ms, *SD* = 7.57) and the 1400 ms (*M* = 42.97 ms, *SD* = 6.74) when compared to the respective onset latencies for baseline trials (*M*_*350*_ = 39.93 ms, *SD* = 6.76, *p* = .038; *M*_*1400*_ = 39.83 ms, *SD* = 6.08, *p* < .001). Figure 3D shows mean OOc EMG latency in baseline and probe trials for each foreperiod length.

The amplitude of OOc EMG was reduced from baseline (*M*_*t-score*_ = 53.02, *SD* = 14.22) in probe trials (*M*_*t-score*_ = 46.72, *SD* = 12.22), as indicated by a significant main effect of trial type, *F*_*(1*, 879.39)_ = 101.66, *p* < .001, *R*^*2*^ = .104. However, the interaction of foreperiod duration with trial type was not statistically significant, *F*_*(2*, 879.69)_ = 2.25, *p* = .106, *R*^*2*^ = .005. Figure 3E shows mean OOc EMG amplitude in baseline and probe trials for each foreperiod duration. Examination of OOc EMG amplitude t-scores as a percentage of baseline across foreperiod lengths indicated a significant main effect of foreperiod duration, *F*_*(2*, 410.88)_ = 6.54, *p* = .002, *R*^*2*^ = .031. Pairwise comparisons indicated the 1400 ms foreperiod length resulted in a significantly greater reduction of OOc EMG amplitudes from baseline to probe in comparison to the 350 ms foreperiod length (*p* = .001) and the 700 ms foreperiod length (*p* = .041; see figure 3F). Finally, we calculated BFs to evaluate the degree of evidence to suggest the OOc response amplitudes after the probe stimulus showed an inhibitory effect. In contrast to corresponding analysis of MEP amplitude in experiment one, OOc responses showed evidence of inhibition for all foreperiod duration conditions. Evidence for an inhibitory effect increased with increasing preparation time, BF_01_ = 24134.85, 5465834, and 223496324928 for the 350 ms, 700 ms, and 1400 ms conditions, respectively.

## Discussion

In this work, we examined whether changes in the magnitude of preparatory suppression can be observed when urgency to prepare a motor response is manipulated, impacting the time available for motor circuits to engage in preparatory inhibition. These manipulations of urgency were achieved by modifying the duration of time that occurs between the start of the movement of a clock hand and the time a prepared motor action should be initiated. We predicted time constraints on preparatory processes would reduce CS suppression and subsequently, in experiment one, examined whether inhibition of the CS tract is modulated by the urgency of preparation. We supplemented this in experiment two by using a LAS to examine whether inhibition of subcortical startle circuits was similarly modulated by the time available for preparation. Intense sensory stimuli are known to trigger prepared actions early and increase the magnitude of movement execution. Therefore, we analysed motor output after the LAS and examined how the effects of a LAS on motor output may also be modulated depending on the urgency of preparation.

### Preparatory inhibition during low-urgency preparation

When urgency to perform a movement was low, in experiment one we observed evidence of the inhibition of CS pathways, as indicated by a reduction of the amplitude of MEPs elicited during motor preparation in comparison to MEPs elicited at a baseline period before the clock sweep began. Interestingly, in experiment two we also observed evidence of an inhibition of subcortical circuits related to the startle reflex, as measured by the latency and amplitude of startle related OOc responses. This is consistent with a recent report of a delay and reduction of magnitude of OOc responses to a LAS presented during preparation and prior to movement onset (Nguyen et al., 2020). Given preparatory suppression was clearly observed for low urgency movements as indexed by both MEPs and startle related OOc EMGs, it may be assumed that during low-urgency preparation, inhibitory effects can affect both cortical and subcortical motor-related circuits. However, there may be multiple inhibitory processes acting on the central nervous system over the course of action preparation. These separate processes may be difficult to discern from one another when there are little time constraints on preparation and the system is able to freely undergo the usual preparatory processes. As such, the examination of inhibition in both the CS tract and subcortical circuits when there is limited time available for preparation can be particularly useful in differentiating these potential processes, as discussed next.

### Preparatory inhibition during time-constrained preparation

There was no evidence of CS suppression when preparation urgency was high in experiment one. In contrast, OOc amplitude after the probe LAS in experiment two was reduced for all foreperiod durations, although the magnitude of suppression was progressively increased as the amount of preparation time was increased. As such, there may be separate processes of premovement suppression which can be observed at different levels of the central nervous system. Therefore, a distinction should be made between preparatory inhibition of the CS tract, and a potentially more global inhibition which can be observed in subcortical circuits. Furthermore, the phenomenon of preparatory inhibition does in fact appear to be modified by the urgency of an impending motor action. In simple and choice RT tasks, preparatory inhibition is known to be sensitive to foreperiod duration. The magnitude of preparatory inhibition has previously been shown to be reduced at longer foreperiod durations (e.g. > 2000 ms) in simple and choice RT tasks (Davranche et al., 2007; Lebon et al., 2016; Touge et al., 1998; Van Elswijk et al., 2007). This is potentially due to impaired temporal estimation of the imperative signal as the foreperiod duration is increased (Jaskowski & Verleger, 1993), limiting the capability of the system to effectively engage preparatory processes. Here we used an anticipatory timing task which maintains high temporal predictability. We have shown an impairment of CS inhibition which appears to occur due to time constraints on preparatory processes. In this context, actions could be initiated on time which suggests CS suppression may not be an obligatory component of preparation.

### Effects of sensory stimulation during time-constrained preparation

#### Sensory stimuli disrupt motor output in the absence of corticospinal suppression

A striking finding was that response vigour was enhanced in probe trials in comparison to control trials when movement urgency was low, but was reduced when movement urgency was high. This indicates that preparatory inhibition may modulate the effects of sensory stimulation on the execution of motor actions. The enhancement of force and vigour observed in the low urgency condition is consistent with previous reports in the StartReact literature, in which a LAS may add neural activity to motor program circuits which results in a greater magnitude of movement execution (Anzak et al., 2011; Marinovic et al., 2015; McInnes et al., 2020; Tresilian & Plooy, 2006; Ulrich et al., 1998). The similar effects on peak force and vigour that occurred in probe trials regardless of whether the probe was a LAS or TMS pulse may be attributable to accessory stimulation induced by the TMS coil discharge, rather than the stimulation provided by TMS itself (Hershenson, 1962). For example, when discharging, the TMS coil produces an auditory click and produces some tactile sensation on the head. Consequently, the TMS coil may have provided bimodal stimulation which contrasts to the unimodal LAS. Bimodal stimuli have been shown to have greater effects on the early triggering of movement and enhancement of vigour in comparison to unimodal stimuli (Marinovic et al., 2015), which may explain the particularly disruptive effects of TMS in the high urgency condition of experiment one. The disruption of vigour that we observed in both of our experiments is similar to that which has been reported by Xu-Wilson et al. (2011), who identified a decrement in the vigour of saccades when TMS was applied shortly before or soon after saccade onset. The effect was observed regardless of the stimulation site over the skull, indicating that like our findings, the observed reduction of vigour could be attributed to the accessory sensory stimulation emitted by the coil discharge. The resemblance of those findings with the disruption of motor output in the absence of CS suppression we observed here may warrant further examination of the manifestation of preparatory inhibition during saccade preparation.

#### Modulation of the orbicularis oculi response with preparation urgency

The latency of OOc responses to the LAS in experiment two mirrored the pattern of inhibition observed in FDI MEPs during experiment one. Suppression of OOc latency was observed during low-urgency preparation but not during high urgency preparation. Amplitude of the OOc response, however, contrasted this. OOc amplitude was suppressed for all foreperiod durations, with an increasing magnitude of suppression with increasing preparation time. One possible explanation for the divergence between OOc latency and amplitude we observed here is the fact that startling stimuli can elicit two separate eyeblink components – the auditory eyeblink reflex and the auditory startle reflex. These can be difficult to distinguish from one another when analysing EMG records. The auditory eyeblink occurs at short latencies and is thought to be mediated by mesencephalic circuits (Brown et al., 1991). This precedes the auditory startle reflex which, when activated, results in a later onset OOc response along with a generalised skeletomotor response. Importantly, the auditory startle reflex is thought to originate from bulbopontine circuits, a pathway of which is distinct from those associated with the auditory eyeblink reflex (Brown et al., 1991). Given the auditory eyeblink is the first component to occur temporally, measurement of OOc onset is most likely to capture this response. Measurement of OOc EMG amplitude on the other hand, may capture the peak of the auditory eyeblink reflex, the auditory startle reflex, or both, depending on which response was largest in a given trial. As such, the distinctive responses of OOc in terms of onset latency versus amplitude may reflect a differing effect of motor preparation on auditory eyeblink versus startle responses. The utility of examining OOc EMG in this context should also be noted. Our findings indicate that the startle response may provide an indication of the inhibition of motor pathways prior to movement initiation without the need for the presentation of electromagnetic stimulation, which may be unsuitable for some participants (Rossi et al., 2009).

### The role of corticospinal suppression during preparation

Overall, the failure to observe evidence of premovement CS suppression in close temporal proximity of movement initiation, when urgency to move was high, brings to question the assumption that this phenomenon is an integral part of motor preparation. Rather, preparatory suppression of the CS tract may reflect a strategy employed by the motor system to protect the prepared reponse from interference. Evaluation of the behavioural effects we observed in contexts where preparatory CS inhibition was evident in comparison to when it was not may shed light on the potential strategic purposes of this phenomenon. The direction of force modulation we observed is opposite to that which we had predicted. We had hypothesised that a lack of preparatory CS inhibition would result in an enhancement of force in probe trials, rather than a reduction. If, as previously proposed, preparatory inhibition serves to keep preparatory activation below initiation threshold (Duque & Ivry, 2009), then we would expect to see effects in line with those that we hypothesised – earlier triggering and enhanced force of movements, as a result of acoustic stimulation close to movement onset, when in the absence of preparatory inhibition. While we observed earlier triggering of movements overall in probe trials, which may again be attributed to the StartReact effect in both of our experiments (Valls-Solé et al., 1999; see also Kohfield, 1971; Nickerson, 1973), the magnitude of the early triggering of movement by the probe stimulus did not appear to differ between foreperiod lengths. The results we present here, and previous findings that preparatory inhibition is present before self-timed movements (Ibáñez et al., 2020), are then inconsistent with the impulse control hypothesis (Duque et al., 2010; Duque & Ivry, 2009).

While we cannot rule out the hypotheses that premovement suppression of the CS tract serves to suppress the initiation of competing response selections or to reduce background noise in motor circuits, our data do not completely fit with these explanations. Therefore, we propose an alternative hypothesis which may better fit the data we report here; preparatory CS inhibition serves as a strategy to protect the prepared movement from external interference. We consider two circumstances which may lead to failures to engage this strategy. First, when there is little time to engage in preparatory processes, the speeded initiation of movement is prioritised which precludes preparatory CS inhibition from taking place, in turn leaving the prepared movement prone to interference from external sources. Alternatively, when the level of motor preparation is low, the motor system may deem it unnecessary to engage in preparatory CS inhibition given there is not a sufficiently prepared movement to be protected from interference, hence leaving movements vulnerable to be disturbed by external sources when they have eventually reached a higher state of preparation closer to the time of initiation. We cannot conclusively rule the absence of CS inhibition, as the temporal location of CS inhibition may simply move to a later time point with increasing urgency. This, however, seems unlikely given CS excitability rise usually occurs 100 ms prior to action onset, leaving less than 150 ms for inhibitory processes to begin and then cease. Furthermore, there was evidence of subcortical suppression, indicating it would be possible for a fast acting inhibition mechanism to manifest at this short timescale. Regardless, there was a direct effect on motor output in the situation when there was no evidence of CS inhibition 250 ms prior to action onset. The finding that time constraints impact CS inhibition to a greater degree than that of startle circuits may support the notion that a more generalised inhibition initially acts on motor circuits which evolves into a more specified inhibition appropriate for the selected action.

## Conclusions

There is a transient suppression of CS excitability prior to movement onset when movement urgency is low and there is sufficient time to prepare a motor response. In contrast, when a movement must be rapidly prepared and initiated and there is little time to engage in preparatory processes, there is no evidence of CS suppression at the same temporal location. Furthermore, both a LAS and TMS pulse were found to impair movement force when presented in the absence of preparatory inhibition. The impairment of force we observed after TMS may be attributed to the accessory stimulation provided by the TMS coil discharge. We conclude that preparatory inhibition may not be a physiologically necessary component of movement preparation, but rather, may reflect a strategy employed by the central nervous system which can serve to protect prepared movements from external interference.

## Acknowledgements

We thank Prof. John Rothwell for his insightful comments on an earlier version of this manuscript. We also thank Thomas Wilkinson and Emily Corti for their assistance with data collection.

